# Lit-OTAR Framework for Extracting Biological Evidences from Literature

**DOI:** 10.1101/2024.03.06.583722

**Authors:** Santosh Tirunagari, Shyamasree Saha, Aravind Venkatesan, Daniel Suveges, Miguel Carmona, Annalisa Buniello, David Ochoa, Johanna McEntyre, Ellen McDonagh, Melissa Harrison

**Affiliations:** Literature Services Team, European Bioinformatics Institute, European Molecular Biology Laboratory (EMBL-EBI), Wellcome Trust Genome Campus, Hinxton, CB10 1SD, Cambridge, United Kingdom; Open Targets, European Bioinformatics Institute, European Molecular Biology Laboratory (EMBL-EBI), Wellcome Trust Genome Campus, Hinxton, CB10 1SD, Cambridge, United Kingdom

**Keywords:** Europe PMC, Open Targets, Bioformer, Deep Learning

## Abstract

The lit-OTAR framework, developed through a collaboration between Europe PMC and Open Targets, leverages deep learning to revolutionise drug discovery by extracting evidence from scientific literature for drug target identification and validation. This novel framework combines Named Entity Recognition (NER) for identifying gene/protein (target), disease, organism, and chemical/drug within scientific texts, and entity normalisation to map these entities to databases like Ensembl, Experimental Factor Ontology (EFO), and ChEMBL. Continuously operational, it has processed over 39 million abstracts and 4.5 million full-text articles and preprints to date, identifying more than 48.5 million unique associations that significantly help accelerate the drug discovery process and scientific research (*>* 29.9m distinct target-disease, 11.8m distinct target-drug and 8.3m distinct disease-drug relationships). The results are made accessible through the Open Targets Platform (https://platform.opentargets.org/) as well as Europe PMC website (SciLite web app) and annotations API (https://europepmc.org/annotationsapi).

## Introduction

The process of identifying drug targets is a critical aspect of drug discovery, requiring an understanding of the molecular and genetic mechanisms of underlying diseases. In this study, a “target” specifically refers to genes or proteins that are investigated for their potential role in disease association and drug discovery. Scientists rely on various sources of evidence such as gene expression changes, genetic variations, and clinical study data to unravel the connections between drugs, targets and diseases (1). To navigate this complexity, the Open Targets Platform (2) was developed as a comprehensive web-based tool that integrates diverse sources of evidence, facilitating the efficient identification of promising drug targets associated with diseases and phenotypes. The Platform combines data from more than 20 different sources to provide target–disease associations, including evidence derived from genetic associations, somatic mutations, known drugs, differential expression, animal models, pathways and systems biology, and text-mining of scientific articles. An integrated score weighs the evidence from each source and type, contributing to an overall score for each target–disease association. This systematic approach harmonises information into a coherent schema and presents it in a user-friendly manner.

Extraction of assertions from scientific articles is an important aspect of this work, and to this end, Europe PMC (3) has played a key supporting role. Europe PMC, a global free biomedical literature repository indexing over 41 million abstracts and 8.7 million full-text articles, provides essential support with its text-mining capabilities. By integrating Europe PMC’s text-mined annotations, the Open Targets Platform harnesses scientific literature as a unique source of information, particularly to identify and elucidate target–disease–drug associations, which are central to its functionality.

The Literature-Open Targets (Lit-OTAR) framework consists of two primary components: Europe PMC text-mining and the Open Targets literature module, as illustrated in Figure 1. Europe PMC utilises deep learning techniques to identify target (gene/protein), disease, and chemical/drug entities within scientific documents. Subsequently, Open Targets performs entity normalisation to accurately map these entities to databases like Ensembl (4), Experimental Factor Ontology (EFO) (5), and ChEMBL (6), while ranking the associations between target–disease–drug mentioned in these documents. The primary goal of this framework is to provide a scalable and continuous service to the scientific community, enabling efficient target validation.

**Fig. 1.**
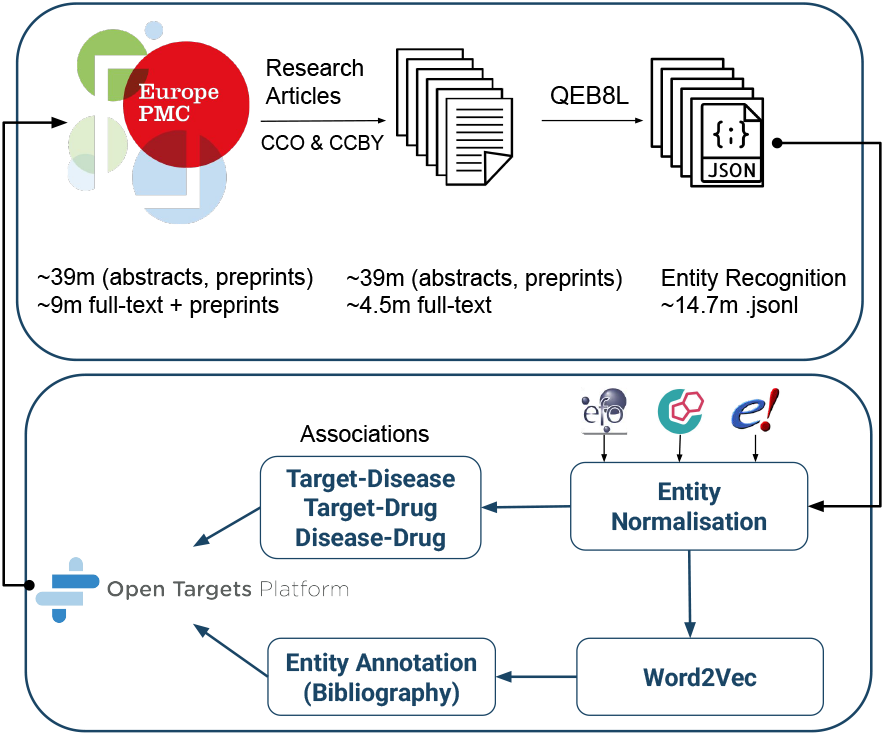
Overview of the Data Selection, Processing, and Accessibility Workflow in Europe PMC and Open Targets Platforms. Refer to Section A (Entity Recognition), Section B (Entity Normalisation) and Section D of the Supplementary Material S7 (Entity Annotation/Bibliography).

Within the existing landscape in biomedical text-mining there are a number of tools focusing on extracting key entities and associations from literature. For instance, DisGeNET (7), SemMedDB (8), LitSense (9), PubTator (10) and PubTator Central (11) provide efficient ways to access high quality text-mined bio-entites. However, these resources are oriented towards access to texmined outputs alone either via highlighting terms or via APIs. The Lit-OTAR work described in here is differently oriented, where the outputs of the framework are mainly integrated with other types of evidences (e.g. RNA expression and pathway analysis) to support systematic identification and prioritisation of therapeutic drug targets in the Open Targets Platform (2, 12). The outputs are highlighted using Europe PMC’s Scilite tool and accessed through the Annotations API.

The Lit-OTAR framework also benefits from an active community that provides documentation, training, and feedback to drive continuous improvements. Community collaboration is facilitated through resources such as the Open Targets Community Portal^1^ and the European PMC’s developer forum^2^, where users share insights and updates.

Our work builds on previous efforts. In our 2017 study (1), we employed dictionary-based methods within a quarterly operational pipeline for data updates, utilising the Europe PMC text-mining pipeline enhanced with custom dictionaries from UniProt and EFO for annotating target and disease names. Although this approach was robust, it faced limitations due to its reliance on manual rules such as abbreviation filters and blacklists of common terms and challenges inherent in biomedical texts. Distinguishing between gene and protein names, spelling variations, and context-specific meanings of abbreviations often led to high recall but low precision (13).

The emergence of modern natural language processing (NLP) techniques (14, 15) has revolutionised text-mining by offering high efficiency and accuracy. Models like BERT (16), BioBERT (17), PubMedBERT (18), and BioFormer (19), trained on extensive biomedical corpora and fine-tuned for specific tasks, have markedly improved the accuracy of entity recognition, managing ambiguities, special characters, acronyms, and identifying synonyms and variations in expression. These advancements not only enhance recall and precision rates but also facilitate the discovery of new biological relationships from the extensive, unstructured data in the life science domain.

In our current study, we have leveraged deep learning techniques, specifically models such as BioBERT and BioFormer, to significantly enhance our pipeline. This updated Lit-OTAR pipeline has been refined to enhance flexibility and modularity, expanding its scope to include a new entity category for chemical/drug. This enhancement has enabled the pipeline to text-mine for associations between drugs and targets, drugs and diseases, in addition to targets and diseases. Furthermore, we have addressed technical challenges such as sentence splitting and boundary detection in complex document structures like tables and figures. The main distinctions between the previous and the current pipeline are detailed in Table T1 of the Supplementary Material.

## Materials and Methods

At the time of writing this article, Europe PMC hosted approximately 39 million journal and preprint abstracts and 9 million full-text journal and preprint articles. However, only a subset of these, specifically 39 million and 4.5 million, respectively, were included due to licensing restrictions (CCO and CC-BY) and their classification as original research articles^3^. This dataset and the subsequent daily addition of the data is run through our custom developed deep learning model for NER extraction (20). The generated output is formatted in JSON, with identified entities treated as matches. Moreover, when two matches or entities occur within the same sentence, they are considered as forming an association or providing evidence. We have completed a study with three experts for treating co-occurrence as association (refer Section Co-occurrence vs Association C). Subsequently, this processed data is forwarded to the Open Targets ETL for the purpose of normalisation (grounding). Disease-related entities are mapped to the Experimental Factor Ontology (EFO), chemical and drug entities to CHEMBL, and gene and protein entities to Ensembl. The resulting data is made accessible through both the Open Targets Platform and Europe PMC annotations APIs, in addition to the Scilite annotations tool (21) on the Europe PMC website (refer to Supplementary Material S7: Section B).

### A. Entity Recognition

To develop deep learning models for the Lit-OTAR framework, we utilised the Europe PMC dataset (13). Initially, this dataset did not include mentions of chemical/drug. To overcome this limitation, we used CHEMDNER BioCreative dataset (22) to annotate the corresponding subset in Europe PMC with chemical/drug mentions, preserving the human-annotated spans. The enriched dataset now covers mentions of gene/protein, disease, chemical/drug, and organism. We trained and evaluated three different models BioBERT, SpaCy (custom trained with PubMed+PMC word2vec (23) over 10 iterations), and Bioformer on this dataset, using the evaluation criteria from SemEval-2013 Task 9.1 (24) (Supplementary Material S2).

### B. Entity Normalisation

The NER tagging of entities occurs at Europe PMC, while normalisation and ranking (Supplementary Material S4) takes place on the Open Targets Platform (Figure 1).

The process involves matching and mapping entities to specific databases/ontologies. The pipeline uses a Word2Vec skip-gram model (25) to transform NER outputs into standardised representations. This includes mapping diseases to the EFO, chemicals to ChEMBL, and genes to Ensembl. The model generates n-dimensional word embeddings, which capture semantic similarities by analysing co-occurrence patterns in the literature.

The model’s calculation of similarity metrics also supports the ranking of entities, determining their relevance to the research context by analysing literature patterns. This method improves the accuracy of entity normalisation and enhances the Open Targets Platform’s value for researchers by offering a comprehensive understanding of biological entity relationships and their potential therapeutic implications (refer to Supplementary Material S3 and Algorithm A1).

### C. Co-occurrence vs Association

A curation task was conducted to annotate 252 sentences for association analysis, with each annotator pair assigned 168 sentences and an intentional overlap of 84 sentences between pairs to measure inter-annotator agreement. The annotation categories for association included the following classes: Altered Expression, Genetic Variation, Regulatory Modification, Any (general or unspecified association), NA (Not Available), and No (No association mentioned). The annotation overlap was measured using Cohen’s Kappa (*K*).

Despite high expectations, the overall Cohen’s Kappa value indicated a variance in perceptions of associations in the range of [0.2 - 0.39], reflecting a low inter-annotator agreement that deemed the overlap unsuitable for machine learning purposes. The association identification presented challenges, evidenced by a low overlap. This difficulty was attributed to various factors, including short sentences lacking clear relations, long sentences with lists of multiple genes/proteins, drugs and diseases, complex sentence structures, and sentences that required additional context for accurate interpretation.

Given the subjectivity in defining associations, we opted to treat co-occurrence as a form of association, including even the absence of explicit associations. This approach allows users to apply post-processing to tailor the data to their specific needs. However, it is important to note that this definition limits the framework’s ability to capture associations that span multiple sentences, such as those involving coreference or inferred context. This constraint may affect the comprehensiveness of extracted associations, as more complex linguistic relationships are challenging to identify.

Following this study, we recognised that association is subjective, leading us to consider co-occurrence as a form of association itself. Consequently, we adjusted our approach to treat co-occurrence as the universal set, acknowledging that any co-occurrence might imply an association, despite the challenges in explicit identification by annotators.

## Results

### D. Entity Recognition

BioBERT led in precision among the models tested, achieving scores of 0.91 (Chemical*/*Drug), 0.90 (Disease), 0.93 (Organism), and 0.91 (Gene*/*Protein), with similarly high recall and F1-scores, demonstrating its effectiveness in entity recognition across various categories. Given the computational demands of BioBERT, our focus shifted towards enhancing the Bioformer-8L model into the QEB8L model. By utilising ONNX for model optimisation, we significantly improved inference speeds without sacrificing performance. Further enhancements through static quantisation not only increased processing speed tenfold but also reduced the model size to approximately 77MB, all while maintaining impressive accuracy with precision scores ranging from 0.85 to 0.94 and F1-scores around 0.88 to 0.89, highlighting its balanced performance as shown in Table 1.

**Table 1.** The performance of different models (Dictionary, BioBERT, spaCy, QEB8L) across four categories (Chemical/Drug, Disease, Organism, Gene*/*Protein) in terms of Precision, Recall, and F1-score metrics evaluated on the Gold Standard test set.

| Category | Model | Precision | Recall | F1-score |
| --- | --- | --- | --- | --- |
| *Chemical/Drug* | Dictionary | 0.53 | 0.34 | 0.41 |
|  | **BioBERT** | **0.91** | **0.92** | **0.92** |
|  | spaCy | 0.80 | 0.73 | 0.76 |
|  | QEB8L | 0.85 | 0.90 | 0.88 |
| *Disease* | Dictionary | 0.48 | 0.74 | 0.58 |
|  | **BioBERT** | **0.90** | 0.80 | 0.85 |
|  | spaCy | 0.82 | 0.71 | 0.76 |
|  | **QEB8L** | **0.90** | **0.88** | **0.89** |
| *Organism* | Dictionary | 0.68 | 0.90 | 0.78 |
|  | **BioBERT** | 0.93 | **0.86** | **0.90** |
|  | spaCy | 0.85 | 0.75 | 0.79 |
|  | **QEB8L** | **0.94** | 0.85 | 0.89 |
| *Gene/Protein* | Dictionary | 0.48 | 0.74 | 0.58 |
|  | **BioBERT** | **0.91** | 0.87 | **0.89** |
|  | spaCy | 0.84 | 0.76 | 0.80 |
|  | **QEB8L** | 0.90 | **0.88** | **0.89** |

SpaCy, recognised for its quick inference speed, presented slightly lower precision scores (0.80 *−* 0.84) and F1-scores (0.76 *−* 0.80) across categories, suggesting its practicality for production-level entity recognition tasks with its efficiency. Conversely, the Dictionary approach (our previous pipeline), while serving as a baseline in the current study achieved lower precision scores (0.48 *−* 0.68) but higher recall in some instances, leading to moderate F1-scores.

Our analysis demonstrated a significant overlap between the gold standard and the QEB8L model, identifying additional entities by the QEB8L model not found in the gold standard. Entities identified by the QEB8L model and dictionaries, but absent in the gold standard, were classified as false positives. Our goal was to minimise these false positives while maximising overlap as illustrated in Figure 2.

**Fig. 2.**
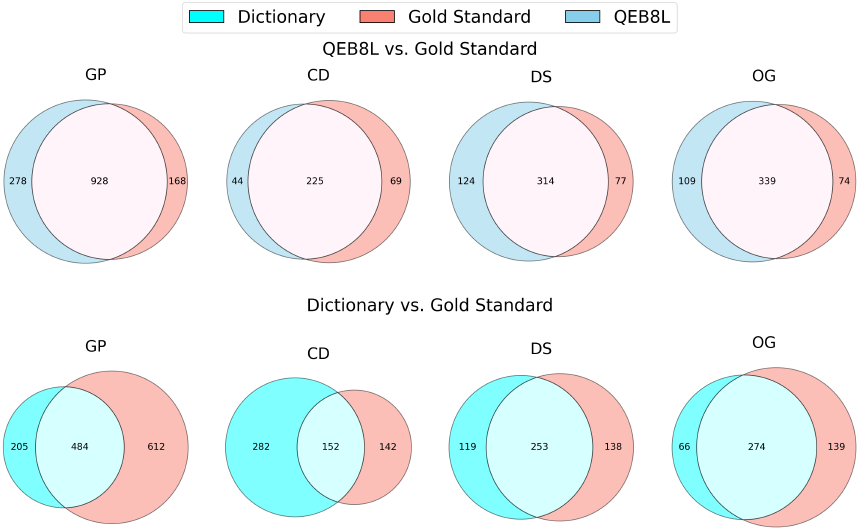
The figure shows the comparison in the number of entity matches between the dictionary-based approach and the gold standard set versus the proposed deep learning approach (QEB8L) to the gold standard (13). The comparison is made between the entities: Gene/Protein (GP), Chemical/Drug (CD), Disease (DS) and Organism (OG) (refer to Supplementary Material S5).

Moving to a deep learning approach, specifically the QEB8L model, was driven by the need to reduce false positives and improve entity coverage. The performance comparison using the gold standard test set demonstrated that the QEB8L model significantly outperformed the previous dictionary-based method, highlighting its advantage. The QEB8L model, trained and evaluated on the gold standard dataset, demonstrated the highest overlap with the gold standard, featuring fewer false positives and false negatives as shown in Figure 2. Some example entities are explained in Supplementary Material S5 (refer to Table T2 for the QEB8L model and Table T3 for the dictionary NER approach).

### E. Entity Normalisation

A large proportion of the recognised entities could be normalised, showing the effectiveness of our methodology in mapping biomedical entities to standardised knowledge bases; Disease to EFO, chemicals to ChEMBL, and Genes to Ensembl. This process is crucial for aggregating and analysing biomedical literature, facilitating the identification of relationships between diseases and potential therapeutic targets.

The entity normalised data, as shown in Table 2, illustrates the scale and complexity of biomedical terminology. Diseases and syndromes alone account for over 220 million entities, with approximately 76.6% successfully normalised to known entities from EFO. However, this represents only 7.6% of unique entity count, highlighting the presence of a long tail of highly heterogeneous and less frequent labels, where rare or variant terms are harder to normalise. The unmapped entities underscore the diversity and complexity of biomedical literature, presenting challenges in achieving complete normalisation, yet remain available for further study^4^.

**Table 2.** Summary of entity recognition and normalisation outcomes across Disease/Syndrome, Chemical/Drug, and Gene/Protein categories. Entity count is the total number of entities identified. Mapped entity count is the number of these entities normalised to a knowledge base. Unique entity count refers to the total distinct entities, while Mapped unique entity count is the subset of those distinct entities that were successfully normalised. Unique mapped references denote the unique knowledge base identifiers to which entities have been mapped.

| Entity type | Entity count | Mapped entity count | Unique entity count | Mapped unique entity count | Unique mapped References |
| --- | --- | --- | --- | --- | --- |
| Disease | 220,392,937 | 168,818,017 (76.6%) | 2,196,439 | 166,497 (7.6%) | 11,561 |
| Chemical/Drug | 122,872,756 | 77,826,420 (63.3%) | 2,213,483 | 76,194 (3.4%) | 10,370 |
| Gene/Protein | 347,835,641 | 197,124,445 (56.7%) | 7,063,573 | 680,368 (9.6%) | 28,778 |

The significant diversity among unnormalised entities necessitates continuous refinement of recognition and normalisation techniques. A literature-based evidence set curated by the Uniprot team was used to benchmark the performance, focusing particularly on disease-to-target associations. This set, comprising of 969 publications with 1, 038 disease–target associations, served as a foundation for evaluating the efficiency of the lit-OTAR pipeline.

The evaluation, detailed in Table 3, demonstrates high match rates for target identification, indicating the Lit-OTAR framework’s potential for mapping biomedical entities to standardised knowledge bases. Conversely, disease recognition and normalisation presented less robust results, highlighting areas for improvement due to the complexity and variability of disease nomenclature.

**Table 3.**
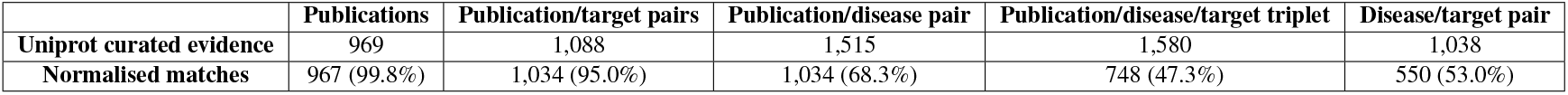
Benchmarking entity recognition and normalisation performance using a UniProt-curated gold standard evidence set.

|  | Publications | Publication/target pairs | Publication/disease pair | Publication/disease/target triplet | Disease/target pair |
| --- | --- | --- | --- | --- | --- |
| Uniprot curated evidence | 969 | 1,088 | 1,515 | 1,580 | 1,038 |
| Normalised matches | 967 (99.8%) | 1,034 (95.0%) | 1,034 (68.3%) | 748 (47.3%) | 550 (53.0%) |

These findings present the strengths and challenges of current Lit-OTAR framework, emphasising the need for advancements in handling the diversity and complexity of disease terms. Interestingly, Lit-OTAR also facilitated unexpected achievements, including the discovery of new disease entities and synonyms (“T2D”), and enhanced data processing capabilities by integrating with databases like EFO and improving analyses with the FDA’s Adverse Events Reporting System (FAERS). The details are presented in Supplementary Material S6.

## Conclusions

The Lit-OTAR framework, a collaboration between Europe PMC and Open Targets, harnesses biomedical literature to advance drug discovery. By applying named entity recognition and entity normalisation, this framework has processed more than 39 million abstracts and 4.5 million full-text articles, identifying around 48.5 million unique associations among target–disease, target–drug, and disease–drug interactions. This study provides insights into the drug discovery process and expands scientific research. In addition, the framework demonstrates the capability to discover new entities and enrich databases and ontologies with previously unrecognised associations. The Lit-OTAR pipeline operates daily, with updates provided quarterly on both the Europe PMC and Open Targets Platforms, ensuring timely access to relevant data for researchers and supporting therapeutic research and development.

## Availability and Implementation

### Data availability

Access the latest data version via FTP^5^, GraphQL API^6^, and Google BigQuery^7^. Further details on the Platforms are presented in the Supplementary Material S7.

### Code availability

The computational frameworks and models supporting this study are distributed across several repositories, maintained by Europe PMC and Open Targets, to ensure broad accessibility and facilitate collaboration.

- The QEB8L model for entity recognition is at https://github.com/ML4LitS/annotation_models.
- The Open Targets daily pipeline, under Europe PMC, is at https://github.com/ML4LitS/otar-maintenance.
- Open Targets’ ETL processes available at https://github.com/opentargets.

## Author contributions statement

All authors contributed significantly. S.T. led writing, NER development, framework productionalisation, and analysis. S.S. set up the daily pipeline and aided experiments. D.S. evaluated normalisation and tested the framework. A.V. and A.B. assisted with writing, while A.V., M.H., and D.O. were pivotal in integrating the framework into Europe PMC. E.M. and J.M. contributed to project ideation and provided key conceptual insights.

## ACKNOWLEDGEMENTS

We thank the curators at Molecular Connections for their biocuration efforts. We would also extend our thanks to Zunaira Shafique and all the development team from Europe PMC and Open Targets Platform for their contributions to the Europe PMC Platform and Open Targets Platform. This work was supported by: the European Molecular Biology Laboratory-European Bioinformatics Institute (S.T, A.V, MH); and the OpenTargets grant 2056 (S.T, S.S).

## Competing interests

No competing interest is declared.

## Supplementary Material

### S1: Old pipeline vs new Lit-OTAR pipeline

**Table T1.**
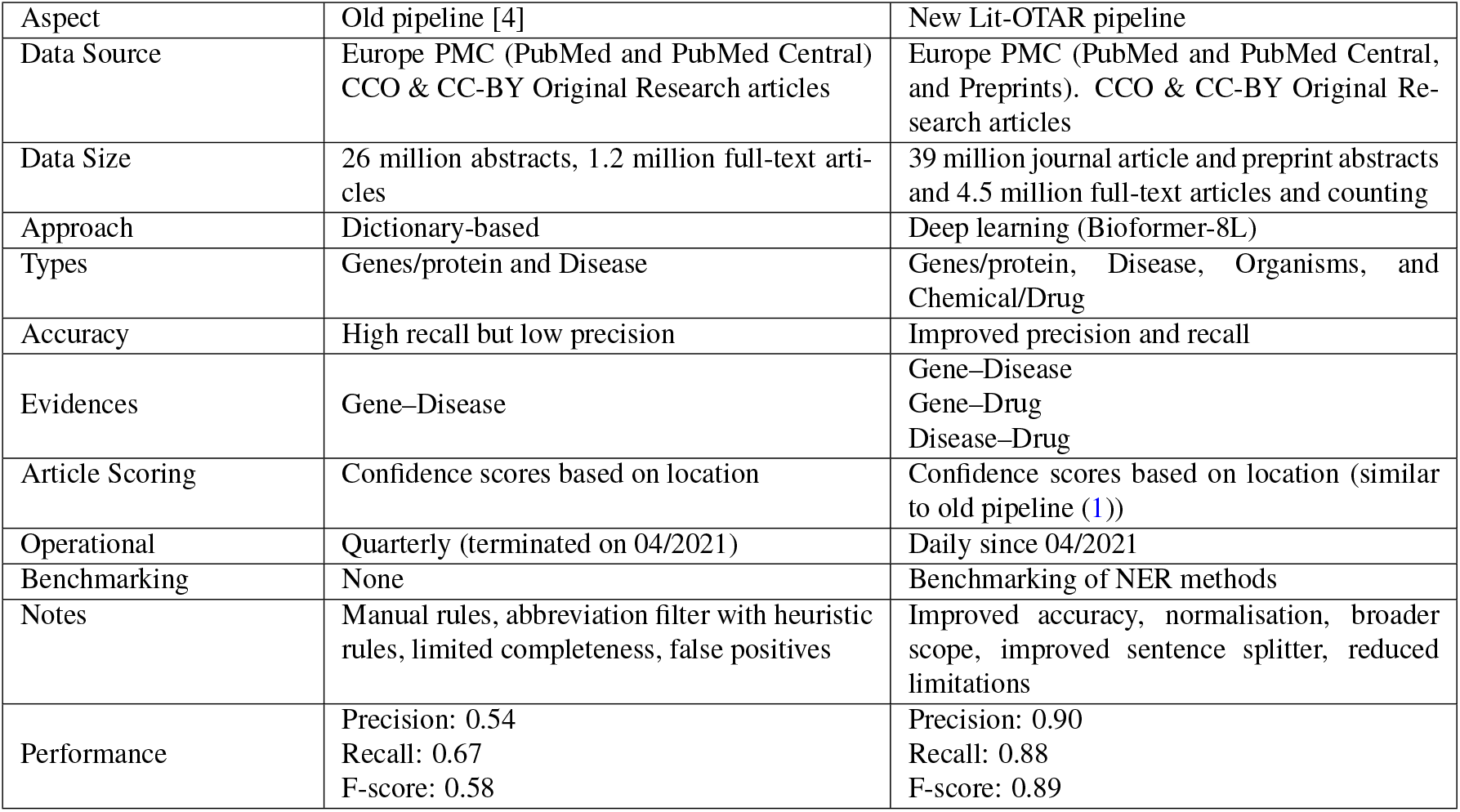
Comparative Analysis of the Old pipeline (1) and the Current pipeline Across Various Aspects

### S2: Evaluation Criteria

The NER was evaluated using the evaluation criteria used by SemEval-2013 Task 9.1 (24), which allows assessment of the system’s performance based on four levels of strictness: strict, exact, partial, and type. Strict evaluation requires both the boundaries and the type of an entity to match exactly with the reference annotation, meaning even slight boundary differences result in a mismatch. Exact evaluation, on the other hand, focuses on the boundaries alone, ensuring they match perfectly without factoring in type accuracy. These levels consider the match of entity boundaries and types. Using these metrics to evaluate the performance of the NER go beyond simple strict classification and take into account partial matching. To compare the differences between the output of the NER system and the correct annotations, two factors were considered: the exact string and the type of the entity. However, because there can be overlapping entities from different categories and data formats, each system per category was evaluated. This means that in certain cases, the counts in the “Strict” and “Exact” cells become equal. Similarly, this applies to the values in the cells that correspond to partial matching and incorrect matching. After evaluating the NER system using the metrics discussed above, the precision, recall, and F1-score was calculated for benchmarking.

The entity-level **precision** and **recall** are computed by deciding when a predicted entity counts as a correct match (COR), in contrast to being labeled as partial (PAR), incorrect (INC), spurious (SPU), or missed (MIS). Once “correct” is defined under a particular matching scheme (Strict, Exact, Partial, or Type), we use the usual formulas^8^:

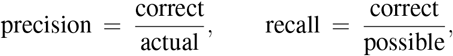

where actual = number of system output entities (TP + FP), possible = number of gold (true) entities (TP + FN).

#### Match Schemes in Detail

1. **Strict**. A system entity is counted as correct (COR) only if:
  - Its boundaries match the gold entity exactly (same start and end tokens),
  - and the type is identical (e.g., both are DISEASE).
  If either boundary or type differs, it is labeled as INC (incorrect), PAR (partial), etc.
2. **Exact**. The boundaries must match exactly, but the entity type is ignored for correctness. Thus, a perfect boundary match is always COR, regardless of the predicted vs. gold type.
3. **Partial**. Any overlap between a system-predicted entity and a gold entity is at least a partial match. Following Batista’s implementation:

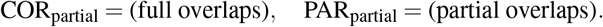

Partial matches contribute half a point in precision and recall:

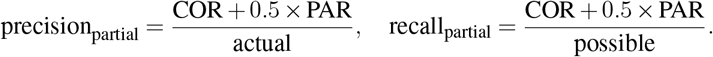

actual Types are ignored in the “Partial” scheme.
4. **Type**. The system entity must overlap with the gold entity and must match its type. Overlaps with the same type are counted as COR (for full overlap) or PAR (for partial). Partial overlaps receive half credit, while overlaps with different types are INC.

Finally, each system entity is ultimately labeled as one of five:

- **COR** – correct
- **INC** – incorrect
- **PAR** – partial match
- **MIS** – missed (the gold entity was not found)
- **SPU** – spurious (the system predicted an entity that does not exist in gold)

Then, for each of the four schemes (Strict, Exact, Partial, Type), we decide what qualifies as “COR.” In the Partial and Type evaluations, partial matches (PAR) count as 0.5 toward precision and recall. Finally, the precision, recall, and F1-score are computed for each of these schemes to benchmark system performance.

### S3: Entity Linking

The entity linking process, implemented in Scala and integrated into the Open Targets pipelines https://github.com/opentargets/platform-etl-backend/blob/master/src/main/scala/io/opentargets/etl/backend/literature/, is designed to efficiently map disease, drug, and gene labels to their corresponding EFO, ChEMBL, or Ensembl identifiers. This approach involves generating a comprehensive lookup table from the Open Targets Platform’s disease, target, and drug indices. Each dataset is processed independently, with all possible labels-such as names, symbols, and retired terms-being expanded for each identifier. To rank the mapped keywords, similarity factor is applied, prioritizing, for example, approved symbols over obsolete names. In cases where a single label maps to multiple identifiers with equal ranking, the process disambiguates by aggregating all related labels and identifiers from the source paper, selecting the most representative identifier based on the assumption that multiple mentions of the same entity in a paper will use various synonyms. This ensures accurate linking and minimizes redundant matches in the evidence.

The Algorithm A1 outlines a structured approach starting with the “Main” Procedure, where the initial data comprising entity matches (named entities), and sentence texts is loaded and passed through various stages. Note that in Europe PMC pipeline, only sentence splitting and section tagging are performed; the sentences are then fed directly to the QEB8L model for entity recognition based on the subword tokenization provided by Bioformer. By contrast, the Open Targets pipeline (Algorithm A1) carries out its own NLP preprocessing.

In the first stage PreprocessData, the data undergoes filtering based on entity types and publication sections. This function also normalizes the text to ensure uniformity (UTF-8 conversion).

Next, the GroundEntities function applies NLP techniques such as tokenization, stopword removal, and stemming. These steps help in breaking down the text into manageable units and prepare the data for entity linking. Following this, the Word2VecModel function trains a Word2Vec model on the preprocessed text, using parameters like window size and iteration count to guide the training process. This trained model serves as the basis for mapping text to entities.

In the MapTextToEntities function, the trained Word2Vec model is applied to map text data to corresponding entities by measuring similarity. A similarity threshold (70%) is employed to determine valid matches, and the function adjusts mappings to optimize performance. Post-processing is handled by the PostProcessOutput function, which resolves co-occurrences and ranks the results based on defined metrics, ensuring that the output is both relevant and accurate.

Finally, the entity mappings, co-occurrence information, and failed mappings are saved to specified output paths (SaveOutput function). This structured pipeline ensures that the NER data is effectively processed, entity linked, and saved for further analysis or reporting.

### S4: Article Scoring and Ranking

The article scoring method in the lit-OTAR framework follows the approach detailed in (1). The algorithm scores scientific articles on their relevance to target–disease associations, helping to rank/prioritise articles by their relevance. The algorithm uses a weighting system that assigns different values to article sections, from full-text articles to abstracts, based on their ability to highlight key entities. For instance, the “Title” section gets the highest weight as it summarises the study’s findings, while the “Introduction” is weighted least, given its focus is on known information. In abstracts, weight is given based on sentence position, with the analysis of 360 MEDLINE abstracts (1) showing that the last sentence, usually detailing results, is considered most significant.

### S5: Examples: False Positives and False Negatives

The examples of entities found through QEB8L but missed in the Gold Standard (False Positives) versus entities found in the Gold Standard but missed through QEB8L (False Negatives) are presented in Table T2. Similarly, Table T3 presents entities found through the Dictionary Approach but missed in the Gold Standard (False Positives) versus entities found in the Gold Standard but missed through the Dictionary Approach (False Negatives).

**Table T2.**
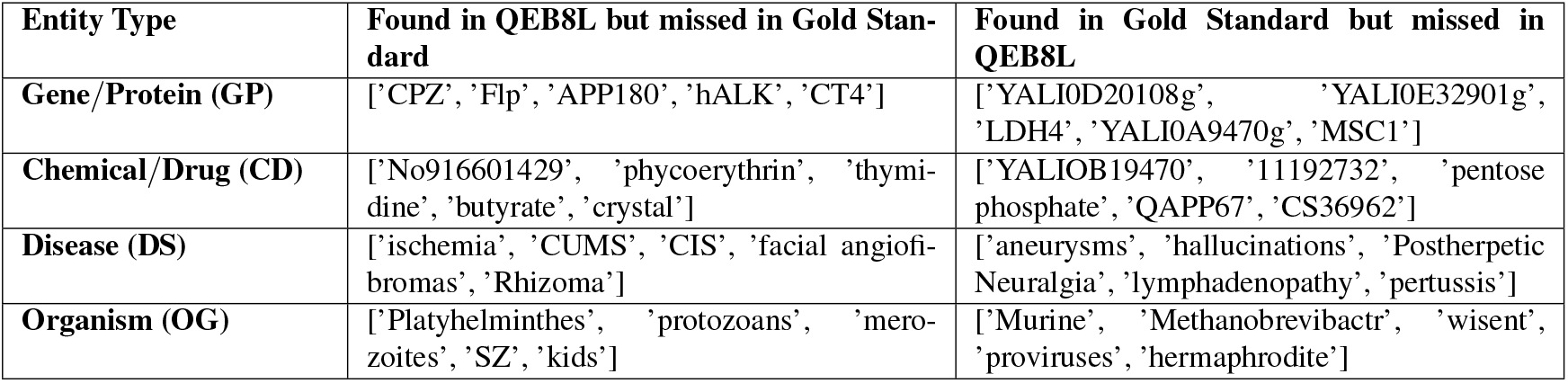
Example Entities Found through QEB8L but Missed in Gold Standard vs. Entities Found in Gold Standard but Missed through QEB8L.

#### Algorithm A1

Entity Linking Pipeline

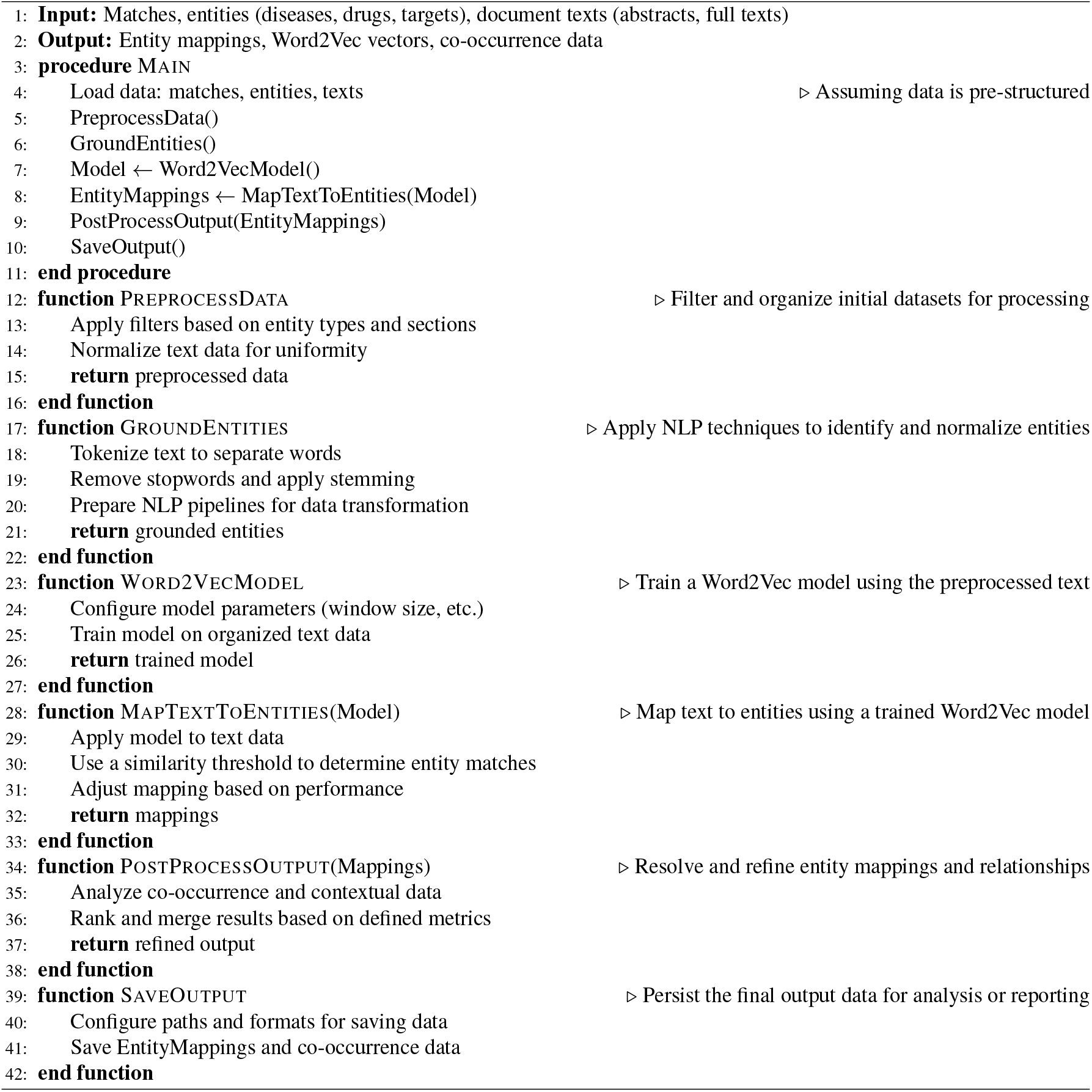

For instance, in Table T3, the organism “cotton” was missed in the Gold Standard. The term “cotton” in the given context refers to bedding material rather than the plant species, as shown in the sentence:

> Each male compartment contained a stainless steel nest-box (130 mm × 130 mm × 130 mm) filled with cotton bedding, a cardboard tube, water bowl, feed tray, and plastic climbing lattice on one wall. (PMCID: PMC4414469, Figure 1)

This differs from its occurrence in the Gold Standard, where “cotton” refers to the plant in an agricultural context:

> Geminiviruses are emerging plant pathogens that infect a wide variety of crops including cotton, cassava, vegetables, ornamental plants, and cereals. (PMCID: PMC3024232, Section Abstract)

Similarly, the chemical entity term “sec” was tagged in one context as referring to “seconds” rather than the intended chemical meaning. Additionally, the term “hermaphrodite” (an organism) was confused with “hermaphroditism”, which may be considered a disorder in certain contexts.

These examples highlight the limitations of the dictionary-based NER approach used to extract entities. While this approach relies on predefined dictionaries and manual rules, such as abbreviation filters and blacklists of common terms, it faces challenges in handling the complexity and variability inherent in biomedical texts. Issues such as distinguishing between gene and protein names (e.g., p53 vs. P53), managing spelling variations (e.g., T2D vs. T2DM), and interpreting context-specific meanings of abbreviations (e.g., AIDS vs. aids) can lead to errors. The example of “cotton,” as discussed earlier, underscores the difficulty in disambiguating context-specific meanings. Additionally, special characters, synonyms, and variations in word choice and sentence structure further complicate entity recognition, often necessitating human interpretation and an exhaustive list of dictionary terms. As a result, this approach, while achieving high recall, often suffers from low precision.

Entities found by QEB8L but missed in the Gold Standard (Table T2) include “merozoites” (PMCID: PMC3097211), which are small, egg-shaped, unicellular organisms that represent a motile stage in the life cycle of malaria parasites:

> After intense multiplication during 2–6 days, depending on the Plasmodium species, mature EEFs release thousands of merozoites, which invade erythrocytes and initiate the pathogenic blood stage cycle.

This was not annotated in the Gold Standard but was correctly identified as an organism by the QEB8L model.

However, there were also instances of misidentification. For example, “Chronic Unpredictable Mild Stress (CUMS)” (PMCID: PMC4931053) was incorrectly identified as a disease, whereas it actually refers to an experimental method. Similarly, in another context (PMCID: PMC5528876), “CIS” (Checklist Individual Strength) was tagged as a disease, likely due to confusion with “clinically isolated syndrome”.

**Table T3.**
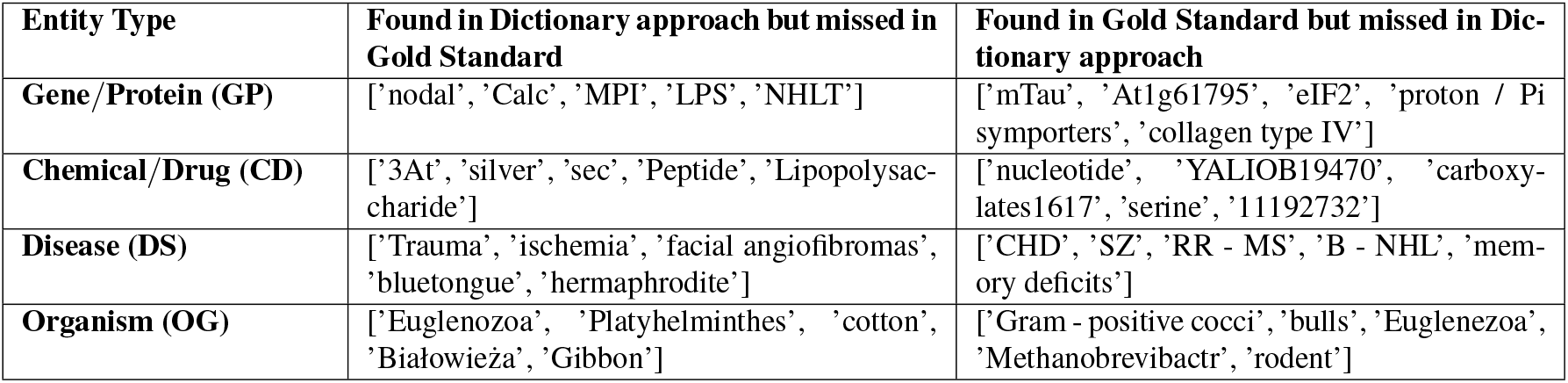
Example Entities Found through Dictionary Approach but Missed in Gold Standard vs. Entities Found in Gold Standard but Missed Through Dictionary Approach.

Given that the QEB8L model tends to tag numerous spurious terms, it is crucial to normalize these terms to a knowledge base using entity linking.

### S6: Other Achievements

The additional entities not found in dictionaries or the gold standard test set (Figure 2), which have been discovered through context learning in deep learning, further facilitate the addition of new entities to databases/ontologies. In one such scenario, the framework has aided in identifying diseases previously unlinked to any specific disease entity within the EFO ^9^. Through a preliminary analysis of frequently occurring non-grounded labels, we identified new synonyms for existing diseases, notably recognising “T2D” as a synonym for Type II Diabetes Mellitus (*EFO*_0001360). This discovery alone added 281,184 matched labels across 29, 040 unique PubMed identifiers (PMIDs), significantly enriching the dataset and enhancing the accuracy of disease-related data mapping.

In another scenario, it enhanced the processing of data from the FDA’s Adverse Events Reporting System (FAERS). This system, which compiles reports of adverse events and medication errors submitted to the FDA, presents unique challenges, such as distinguishing between a drug’s adverse events and its indications. To address this, an increase in EFO cross-references to MedDRA was necessary in order to find out whether excluding reports where the adverse event matches the drug indication could improve the analytical outcomes’ power and effectiveness. The normalisation pipeline developed for the Lit-OTAR was able to map a significant portion of MedDRA^10^ labels associated with adverse reactions to their corresponding EFO terms. This effort resulted in a cross-reference list containing approximately 10,000 mappings, which are assessed to be highly reliable.

### S7: Data Platforms

The datasets generated are made available through both Open Targets Platform and Europe PMC. While Open Targets Platform provides a web interface for data exploration and as bulk download, in Europe PMC the datasets are accessible both via the Annotations API and the website.

#### A. Europe PMC Annotations API

The Europe PMC Annotations API^11^ is one of the main methods of accessing texted mined outputs (also called annotations) hosted by Europe PMC. Derived from both abstracts and open access full-text articles, these annotations are an invaluable resource for researchers needing programmatic access to the vast repository. One of the motivations of this is to make text-mined annotations available to the larger scientific community. To this end, the annotations are modelled based on the W3C Web Annotation Data Model^12^. This has ensured the annotations are standardised and, most importantly, FAIRified for wider consumption. The annotations are made available under the Apache License Version 2.0^13^.

**Fig. F1.**
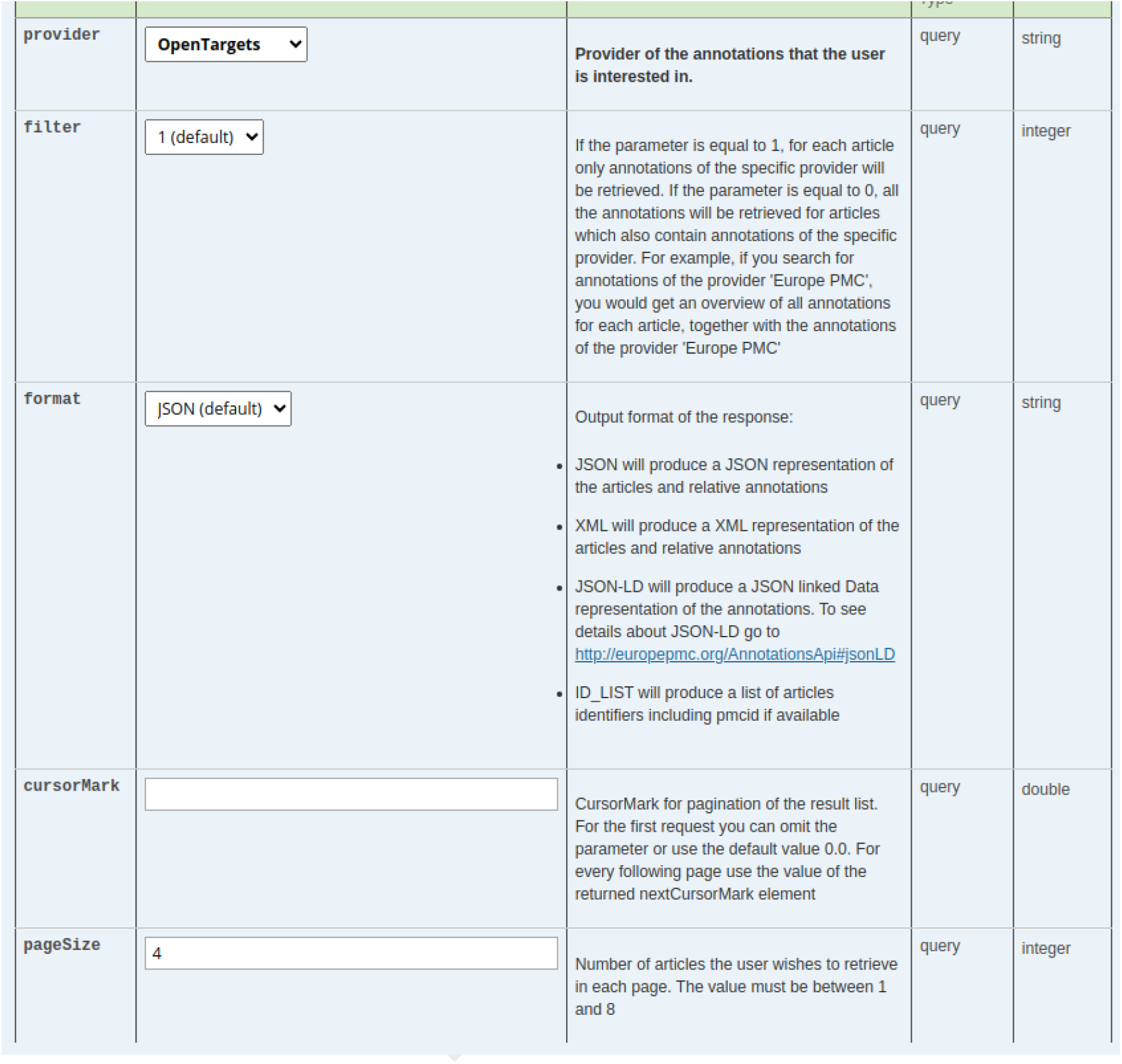
Accessing annotations by OpenTargets on the Europe PMC annotations API

The API’s RESTful architecture offers a modular structure, facilitating various functionalities essential for fetching specific annotations based on article IDs, entities, providers, relationships, or article sections. This flexibility is crucial for researchers aiming to extract detailed and targeted information from the literature. The functionality varies from fetching annotations by article, entity name and provider (e.g. OpenTargets) to relationships such as gene-disease relationships (see Figure F1).

The API delivers results in various formats, including JSON, XML, and ID_LIST (for article identifiers), catering to different user preferences and requirements. Moreover, the annotations are available in JSONLD format, providing a graph representation that enhances data interoperability and integration.

This extensive accessibility to annotated biomedical literature through the Europe PMC Annotations API significantly empowers researchers, pharmaceutical companies facilitating the extraction and analysis of rich datasets for advancing scientific discoveries.

#### B. Visualisations on Europe PMC Website

SciLite (21) is an annotation tool integrated into Europe PMC that highlights key biological concepts within scientific articles. SciLite enables researchers to quickly grasp the important elements of a paper, facilitating more efficient data discovery and making it easier to cross-reference information. The application makes API requests using the Annotations API to fetch all relevant annotations for a given article (see Figure F3). A detailed description of the design behind SciLite can be found here.^14^ SciLite is one of the infrastructural components of the Europe PMC annotation platform. The platform is open for text-mined outputs from any source to be shared and displayed seamlessly on content. This is enabled by the use of the (W3C recommended) Web Annotation Data Model http://www.w3.org/TR/annotation-model/. This aspect differentiates SciLite from other tools such as PubTator (10).

**Fig. F2.**
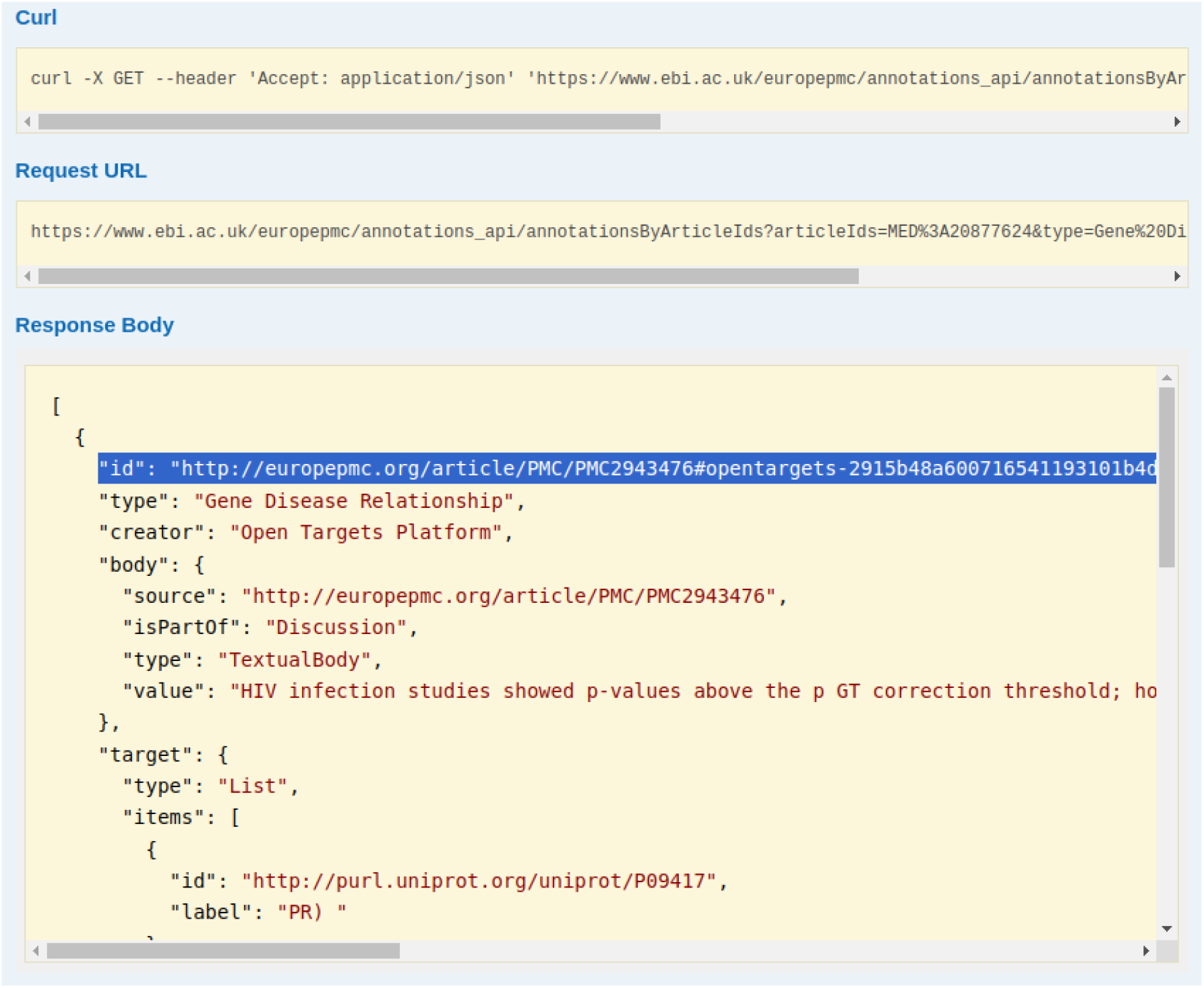
The screenshot of the Open Target annotation (in JSON-LD format) retrieved from the Europe PMC Annotations API for the article with PMID 20877624. The annotation URI for the Open Target annotation is highlighted.

#### C. Making annotations FAIR and Citable

One of the main aims of establishing Europe PMC Annotation API is to make text-mined annotations FAIR, where users are able cite the text-mined entities and relationships. This allows the consumer of the content to understand and trace the source of information. The Web Annotation Model specification allows annotations to be uniquely identified using URIs, offering a mechanism to cite annotations across multiple Platforms. To this end, for all Open Targets annotations we have minted resolvable annotation URIs. For instance, for a given article ID the corresponding Open Target annotation can be retrieved in JSON-LD format, that will contain the annotation URI (see Figure F2).

**Fig. F3.**
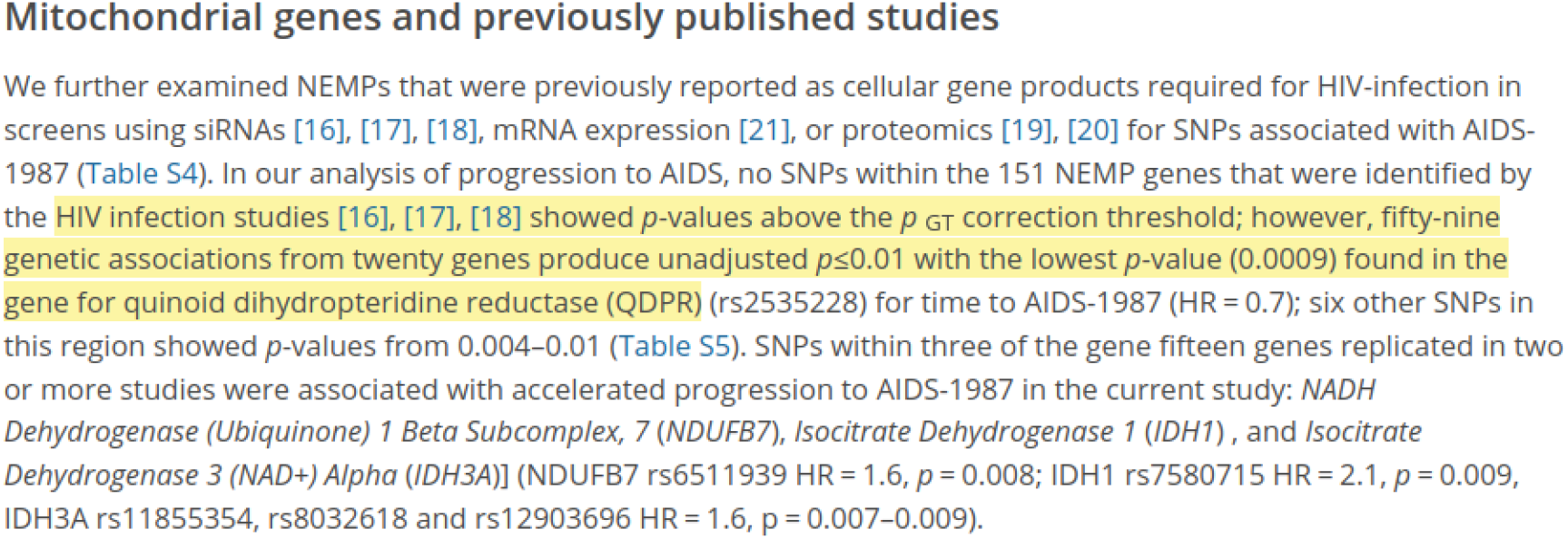
SciLite annotation tool highlighting the gene-disease association between HIV infection and QDPR using the LinkBack call. The LinkBack feature is based on the LinkBack API (which accepts the unique ‘code’ in an annotation ID) and text-annotator.

#### D. Results in the Open Targets Platform

The Europe PMC dataset is utilised in two ways: The first is to use the dataset to extract evidence for the association of targets and diseases. Co-occurrences of target and disease entities are considered evidence for the association of those entities. In detail, when a target and a disease are mentioned in the same sentence within a publication, this constitutes one piece of Europe PMC evidence for the association of that target and that disease [Figure F4 (a)]. The second involves using the dataset to provide context to the Platform entities. Users can browse the available literature for the entity of their choice through the Bibliography widget, for example all the papers linked to cystic fibrosis [Figure F4(b)]. For more details refer to article (2, 12).

### S8: Comparison with Other Tools

The outputs from the lit-OTAR framework are visualised and accessed through Europe PMC’s Scilite/Annotations API and Open Targets Platform. Table T4 present comparison of various tools including SemMedDB (8), LitSense (9), PubTator (10) and PubTator Central (11) which provide similar text-mining outputs.

**Fig. F4.**
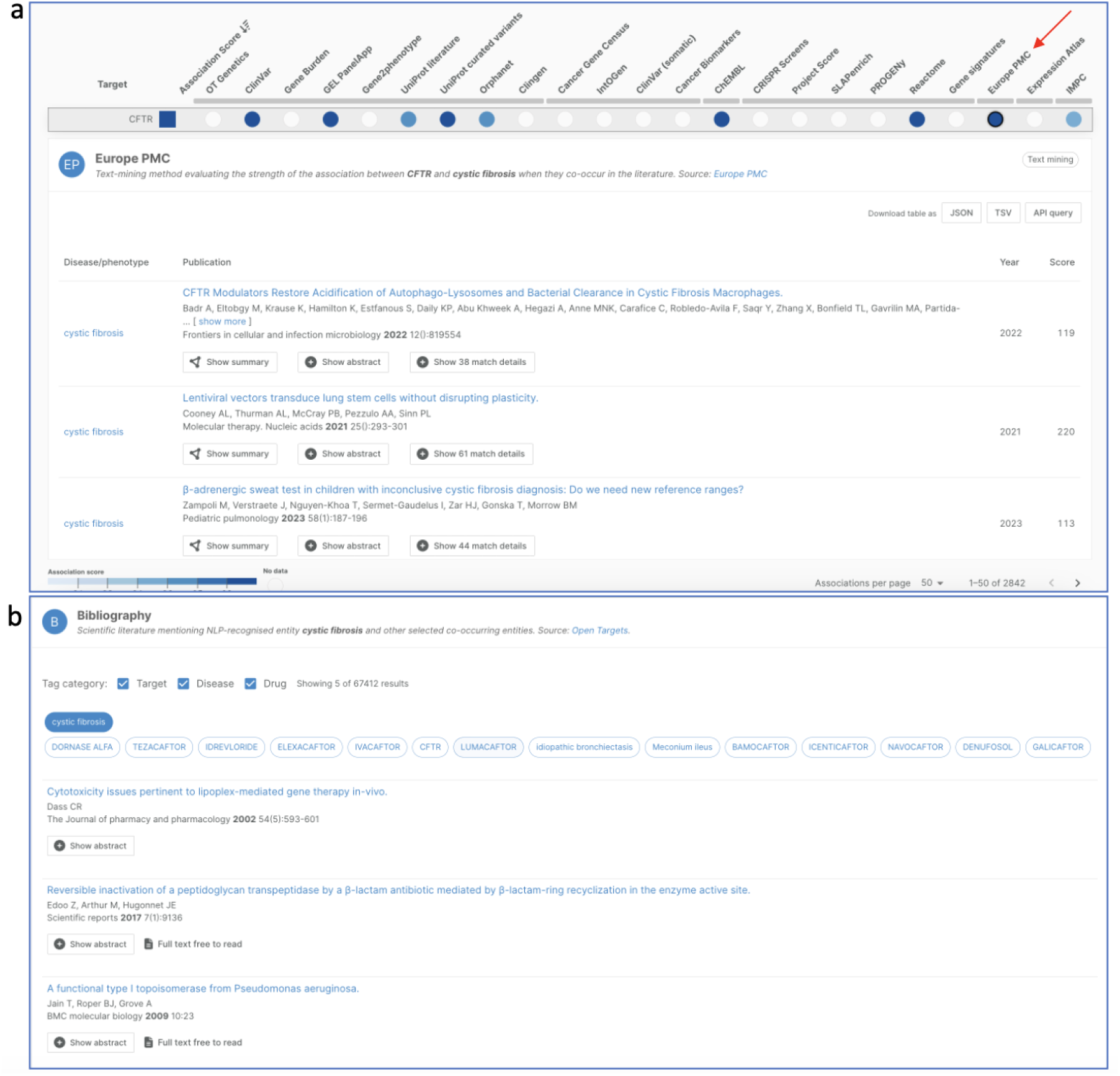
Summary of how the lit-OTAR results are utilised and visualised in the Open Targets Platform. a. Europe PMC (red arrow) as data source for evidence of target–disease associations; b. Bibliography widget from a disease profile page.

**Table T4.**
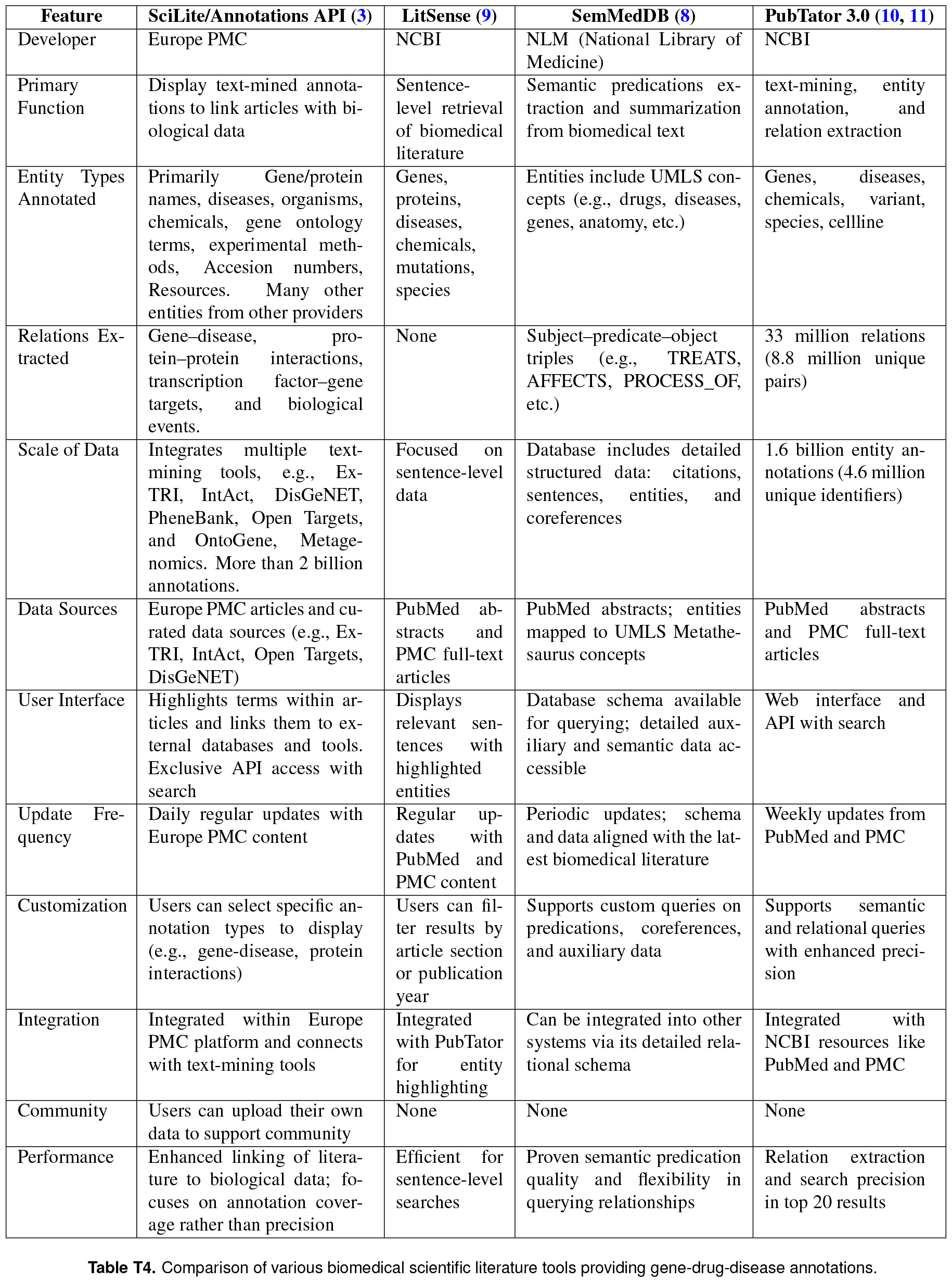
Comparison of various biomedical scientific literature tools providing gene-drug-disease annotations.

## Footnotes

1 https://community.opentargets.org

2 https://groups.google.com/a/ebi.ac.uk/g/epmc-webservices

3 https://europepmc.org/Copyright

4 https://ftp.ebi.ac.uk/pub/databases/opentargets/platform/latest/output/etl/json/literature/failedCooccurrences/

5 https://ftp.ebi.ac.uk/pub/databases/opentargets/platform/latest/output/etl/json/literature/

6 https://api.platform.opentargets.org/api/v4/graphql/browser

7 https://platform-docs.opentargets.org/data-access/google-bigquery

8 http://www.davidsbatista.net/blog/2018/05/09/Named_Entity_Evaluation/

9 https://github.com/opentargets/issues/issues/1555?ref=blog.opentargets.org?ref=blog.opentargets.org

10 https://www.meddra.org/?ref=blog.opentargets.org

11 www.europepmc.org/AnnotationsApi

12 https://www.w3.org/TR/annotation-model/

13 https://www.apache.org/licenses/LICENSE-2.0

14 https://europepmc.github.io/techblog/algorithm/2018/07/04/locating-text-html-pages.html

## Notes

### Competing Interest Statement

The authors have declared no competing interest.

### Summary of Updates

Thank you very much to the reviewers for taking the time to provide us with detailed feedback. We greatly appreciate your comments and have carefully considered each point raised to enhance the quality of this paper. We would also like to extend our sincere gratitude to the Associate Editor for their valuable guidance throughout the review process. New text has been highlighted in purple (in the revised version). Table 3 has been transposed to address spacing issues. Three footnote URLs have been replaced with proper citations for the corresponding knowledge bases (EFO, ChEMBL, and Ensembl). We have reworked the Introduction section to remove grammatical errors and improve readability. Responses to the individual points raised by the reviewers are provided below.

