## Supplementary Material: for "Lit-OTAR Framework for Extracting Biological Evidences from Literature"

### S1: Old pipeline vs new Lit-OTAR pipeline

| Aspect | Old pipeline [4] | New Lit-OTAR pipeline |
|---|---|---|
| Data Source | Europe PMC (PubMed and PubMed Central) CCO & CC-BY Original Research articles | Europe PMC (PubMed and PubMed Central, and Preprints). CCO & CC-BY Original Research articles |
| Data Size | 26 million abstracts, 1.2 million full-text articles | 39 million journal article and preprint abstracts and 4.5 million full-text articles and counting |
| Approach | Dictionary-based | Deep learning (Bioformer-8L) |
| Types | Genes/protein and Disease | Genes/protein, Disease, Organisms, and Chemical/Drug |
| Accuracy | High recall but low precision | Improved precision and recall |
| Evidences | Gene–Disease | Gene–Disease<br>Gene–Drug<br>Disease–Drug |
| Article Scoring | Confidence scores based on location | Confidence scores based on location (similar to old pipeline (1)) |
| Operational | Quarterly (terminated on 04/2021) | Daily since 04/2021 |
| Benchmarking | None | Benchmarking of NER methods |
| Notes | Manual rules, abbreviation filter with heuristic rules, limited completeness, false positives | Improved accuracy, normalisation, broader scope, improved sentence splitter, reduced limitations |
| Performance | Precision: 0.54<br>Recall: 0.67<br>F-score: 0.58 | Precision: 0.90<br>Recall: 0.88<br>F-score: 0.89 |

**Table T1.** Comparative Analysis of the Old pipeline (1) and the Current pipeline Across Various Aspects

DRAFT

### S2: Evaluation Criteria

The NER was evaluated using the evaluation criteria used by SemEval-2013 Task 9.1 (24), which allows assessment of the system's performance based on four levels of strictness: strict, exact, partial, and type. Strict evaluation requires both the boundaries and the type of an entity to match exactly with the reference annotation, meaning even slight boundary differences result in a mismatch. Exact evaluation, on the other hand, focuses on the boundaries alone, ensuring they match perfectly without factoring in type accuracy. These levels consider the match of entity boundaries and types. Using these metrics to evaluate the performance of the NER go beyond simple strict classification and take into account partial matching. To compare the differences between the output of the NER system and the correct annotations, two factors were considered: the exact string and the type of the entity. However, because there can be overlapping entities from different categories and data formats, each system per category was evaluated. This means that in certain cases, the counts in the "Strict" and "Exact" cells become equal. Similarly, this applies to the values in the cells that correspond to partial matching and incorrect matching. After evaluating the NER system using the metrics discussed above, the precision, recall, and F1-score was calculated for benchmarking.

The entity-level **precision** and **recall** are computed by deciding when a predicted entity counts as a correct match (COR), in contrast to being labeled as partial (PAR), incorrect (INC), spurious (SPU), or missed (MIS). Once "correct" is defined under a particular matching scheme (Strict, Exact, Partial, or Type), we use the usual formulas[8]:

$$\text{precision} = \frac{\text{correct}}{\text{actual}}, \qquad \text{recall} = \frac{\text{correct}}{\text{possible}},$$

where

$$\text{actual} = \text{number of system output entities (TP + FP)}, \quad \text{possible} = \text{number of gold (true) entities (TP + FN)}.$$

#### Match Schemes in Detail.

1. **Strict.** A system entity is counted as correct (COR) only if:
   - Its boundaries match the gold entity exactly (same start and end tokens),
   - and the type is identical (e.g., both are `DISEASE`).

   If either boundary or type differs, it is labeled as INC (incorrect), PAR (partial), etc.
2. **Exact.** The boundaries must match exactly, but the entity type is ignored for correctness. Thus, a perfect boundary match is always COR, regardless of the predicted vs. gold type.
3. **Partial.** Any overlap between a system-predicted entity and a gold entity is at least a partial match. Following Batista's implementation:

   $$\text{COR}_{\text{partial}} = \text{(full overlaps)}, \quad \text{PAR}_{\text{partial}} = \text{(partial overlaps)}.$$

   Partial matches contribute half a point in precision and recall:

   $$\text{precision}_{\text{partial}} = \frac{\text{COR} + 0.5 \times \text{PAR}}{\text{actual}}, \quad \text{recall}_{\text{partial}} = \frac{\text{COR} + 0.5 \times \text{PAR}}{\text{possible}}.$$

**Algorithm A1** Entity Linking Pipeline

```
Input: Matches, entities (diseases, drugs, targets), document texts (abstracts, full texts)
Output: Entity mappings, Word2Vec vectors, co-occurrence data
procedure Main
    Load data: matches, entities, texts                    ▷ Assuming data is pre-structured
    PreprocessData()
    GroundEntities()
    Model ← Word2VecModel()
    EntityMappings ← MapTextToEntities(Model)
    PostProcessOutput(EntityMappings)
    SaveOutput()
end procedure
function PreprocessData                    ▷ Filter and organize initial datasets for processing
    Apply filters based on entity types and sections
    Normalize text data for uniformity
    return preprocessed data
end function
function GroundEntities                    ▷ Apply NLP techniques to identify and normalize entities
    Tokenize text to separate words
    Remove stopwords and apply stemming
    Prepare NLP pipelines for data transformation
    return grounded entities
end function
function Word2VecModel                    ▷ Train a Word2Vec model using the preprocessed text
    Configure model parameters (window size, etc.)
    Train model on organized text data
    return trained model
end function
function MapTextToEntities(Model)                    ▷ Map text to entities using a trained Word2Vec model
    Apply model to text data
    Use a similarity threshold to determine entity matches
    Adjust mapping based on performance
    return mappings
end function
function PostProcessOutput(Mappings)                    ▷ Resolve and refine entity mappings and relationships
    Analyze co-occurrence and contextual data
    Rank and merge results based on defined metrics
    return refined output
end function
function SaveOutput                    ▷ Persist the final output data for analysis or reporting
    Configure paths and formats for saving data
    Save EntityMappings and co-occurrence data
end function
```

DRAFT

Table T2. Similarly, Table T3 presents entities found through the Dictionary Approach but missed in the Gold Standard (False Positives) versus entities found in the Gold Standard but missed through the Dictionary Approach (False Negatives).

| **Entity Type** | **Found in QEB8L but missed in Gold Standard** | **Found in Gold Standard but missed in QEB8L** |
|---|---|---|
| **Gene/Protein (GP)** | [’CPZ’, ’Flp’, ’APP180’, ’hALK’, ’CT4’] | [’YALI0D20108g’, ’YALI0E32901g’, ’LDH4’, ’YALI0A9470g’, ’MSC1’] |
| **Chemical/Drug (CD)** | [’No916601429’, ’phycoerythrin’, ’thymidine’, ’butyrate’, ’crystal’] | [’YALIOB19470’, ’11192732’, ’pentose phosphate’, ’QAPP67’, ’CS36962’] |
| **Disease (DS)** | [’ischemia’, ’CUMS’, ’CIS’, ’facial angiofibromas’, ’Rhizoma’] | [’aneurysms’, ’hallucinations’, ’Postherpetic Neuralgia’, ’lymphadenopathy’, ’pertussis’] |
| **Organism (OG)** | [’Platyhelminthes’, ’protozoans’, ’merozoites’, ’SZ’, ’kids’] | [’Murine’, ’Methanobrevibactr’, ’wisent’, ’proviruses’, ’hermaphrodite’] |

| Entity Type | Found in Dictionary approach but missed in Gold Standard | Found in Gold Standard but missed in Dictionary approach |
|---|---|---|
| **Gene/Protein (GP)** | ['nodal', 'Calc', 'MPI', 'LPS', 'NHLT'] | ['mTau', 'At1g61795', 'eIF2', 'proton / Pi symporters', 'collagen type IV'] |
| **Chemical/Drug (CD)** | ['3At', 'silver', 'sec', 'Peptide', 'Lipopolysaccharide'] | ['nucleotide', 'YALIOB19470', 'carboxylates1617', 'serine', '11192732'] |
| **Disease (DS)** | ['Trauma', 'ischemia', 'facial angiofibromas', 'bluetongue', 'hermaphrodite'] | ['CHD', 'SZ', 'RR - MS', 'B - NHL', 'memory deficits'] |
| **Organism (OG)** | ['Euglenozoa', 'Platyhelminthes', 'cotton', 'Białowieża', 'Gibbon'] | ['Gram - positive cocci', 'bulls', 'Euglenezoa', 'Methanobrevibactr', 'rodent'] |

[9] https://github.com/opentargets/issues/issues/1555?ref=blog.opentargets.org?ref=blog.opentargets.org
[10] https://www.meddra.org/?ref=blog.opentargets.org
[11] www.europepmc.org/AnnotationsApi

DRAFT

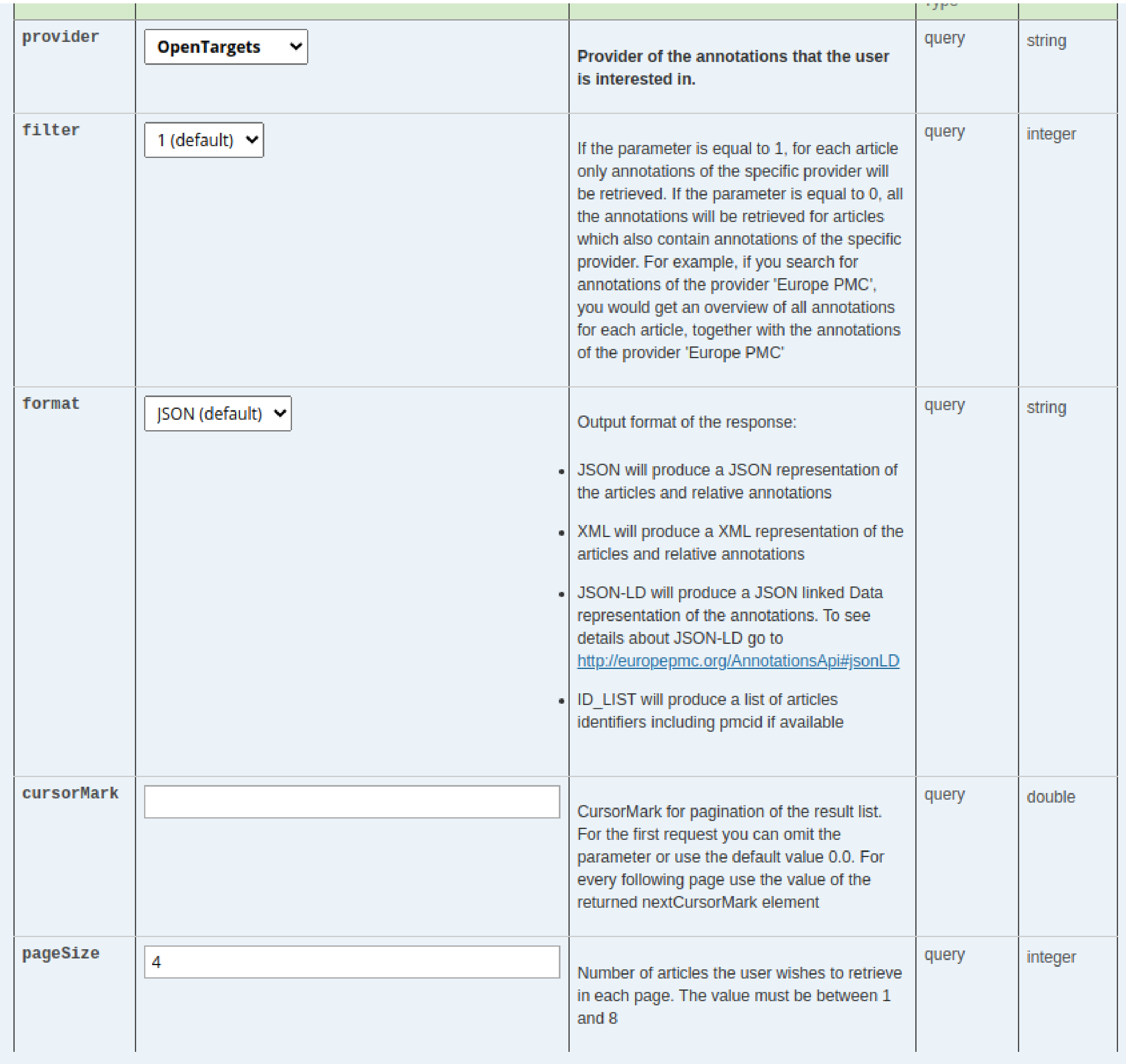

| | | | | |
|---|---|---|---|---|
| provider | OpenTargets | Provider of the annotations that the user is interested in. | query | string |
| filter | 1 (default) | If the parameter is equal to 1, for each article only annotations of the specific provider will be retrieved. If the parameter is equal to 0, all the annotations will be retrieved for articles which also contain annotations of the specific provider. For example, if you search for annotations of the provider 'Europe PMC', you would get an overview of all annotations for each article, together with the annotations of the provider 'Europe PMC' | query | integer |
| format | JSON (default) | Output format of the response: • JSON will produce a JSON representation of the articles and relative annotations • XML will produce a XML representation of the articles and relative annotations • JSON-LD will produce a JSON linked Data representation of the annotations. To see details about JSON-LD go to http://europepmc.org/AnnotationsApi#jsonLD • ID_LIST will produce a list of articles identifiers including pmcid if available | query | string |
| cursorMark | | CursorMark for pagination of the result list. For the first request you can omit the parameter or use the default value 0.0. For every following page use the value of the returned nextCursorMark element | query | double |
| pageSize | 4 | Number of articles the user wishes to retrieve in each page. The value must be between 1 and 8 | query | integer |

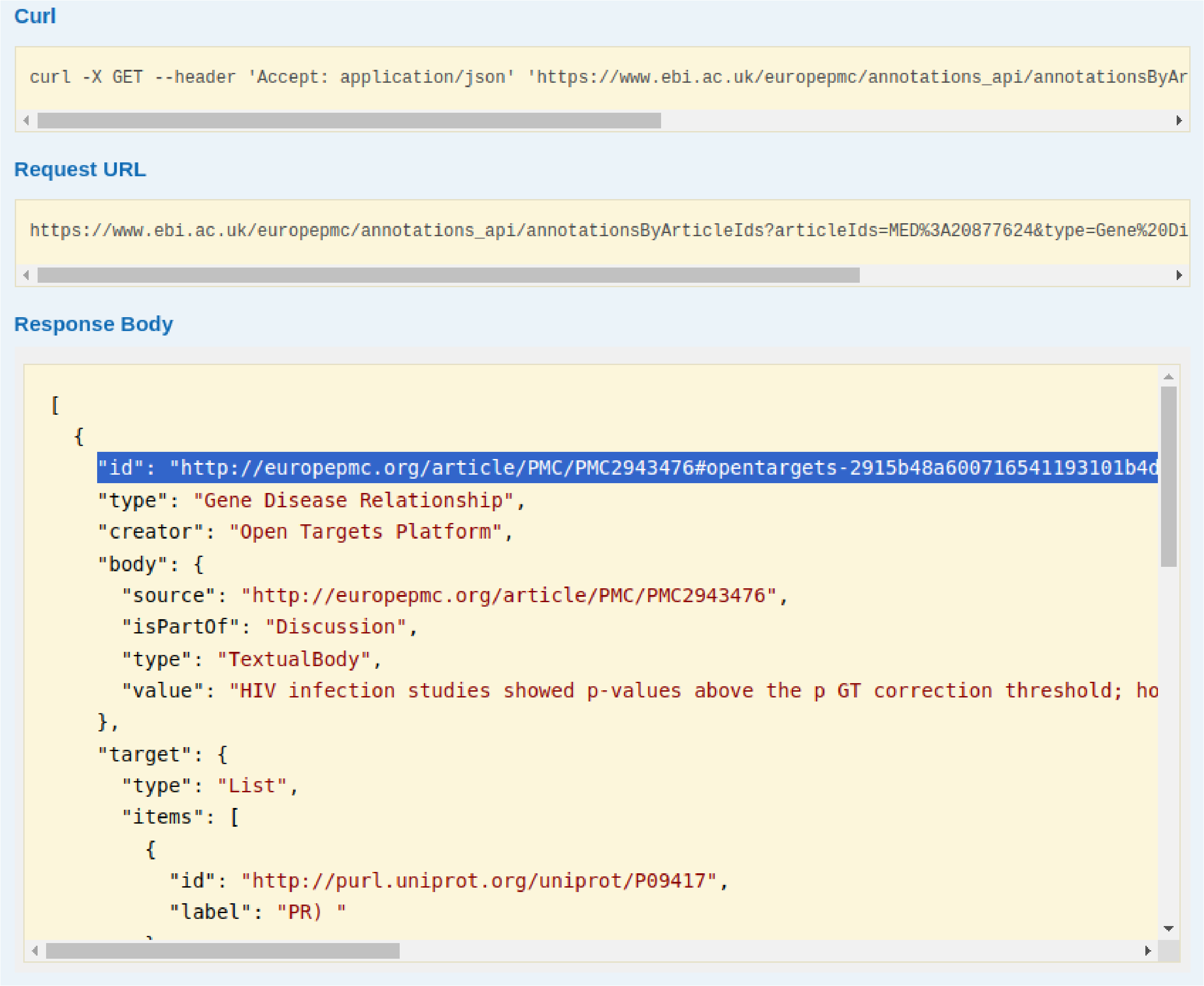


**Fig. F2.** The screenshot of the Open Target annotation (in JSON-LD format) retrieved from the Europe PMC Annotations API for the article with PMID 20877624. The annotation URI for the Open Target annotation is highlighted.

[14] `https://europepmc.github.io/techblog/algorithm/2018/07/04/locating-text-html-pages.html`

**C. Making annotations FAIR and Citable.** One of the main aims of establishing Europe PMC Annotation API is to make text-mined annotations FAIR, where users are able cite the text-mined entities and relationships. This allows the consumer of the content to understand and trace the source of information. The Web Annotation Model specification allows annotations to be uniquely identified using URIs, offering a mechanism to cite annotations across multiple Platforms. To this end, for all Open Targets annotations we have minted resolvable annotation URIs. For instance, for a given article ID the corresponding Open Target annotation can be retrieved in JSON-LD format, that will contain the annotation URI (see Figure F2).

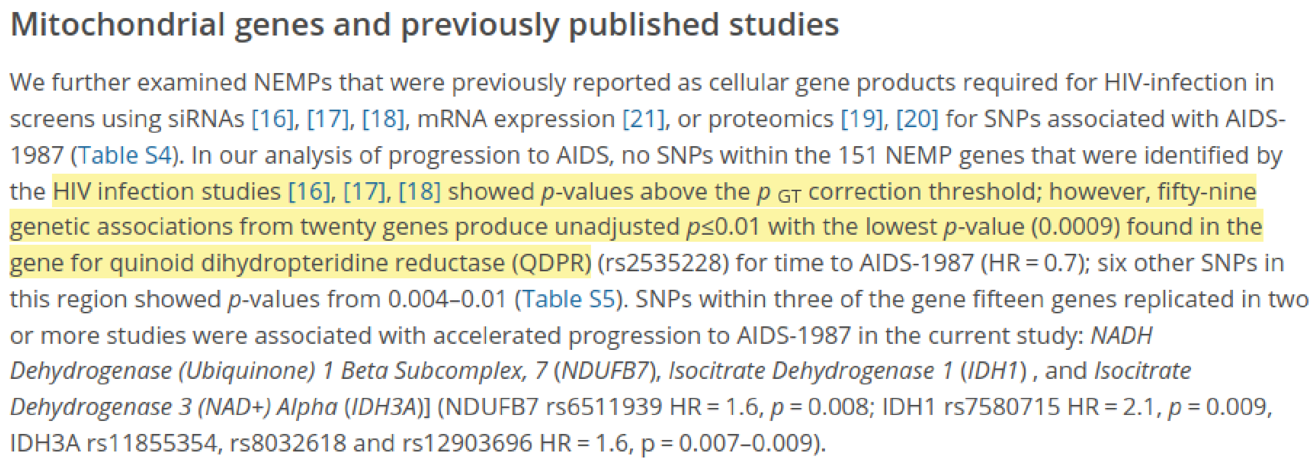

### Mitochondrial genes and previously published studies

We further examined NEMPs that were previously reported as cellular gene products required for HIV-infection in screens using siRNAs [16], [17], [18], mRNA expression [21], or proteomics [19], [20] for SNPs associated with AIDS-1987 (Table S4). In our analysis of progression to AIDS, no SNPs within the 151 NEMP genes that were identified by the HIV infection studies [16], [17], [18] showed *p*-values above the $p_{GT}$ correction threshold; however, fifty-nine genetic associations from twenty genes produce unadjusted $p \leq 0.01$ with the lowest *p*-value (0.0009) found in the gene for quinoid dihydropteridine reductase (QDPR) (rs2535228) for time to AIDS-1987 (HR = 0.7); six other SNPs in this region showed *p*-values from 0.004–0.01 (Table S5). SNPs within three of the gene fifteen genes replicated in two or more studies were associated with accelerated progression to AIDS-1987 in the current study: *NADH Dehydrogenase (Ubiquinone) 1 Beta Subcomplex, 7* (*NDUFB7*), *Isocitrate Dehydrogenase 1* (*IDH1*) , and *Isocitrate Dehydrogenase 3 (NAD+) Alpha* (*IDH3A*)] (NDUFB7 rs6511939 HR = 1.6, $p = 0.008$; IDH1 rs7580715 HR = 2.1, $p = 0.009$, IDH3A rs11855354, rs8032618 and rs12903696 HR = 1.6, p = 0.007–0.009).

**Fig. F3.** SciLite annotation tool highlighting the gene-disease association between HIV infection and QDPR using the LinkBack call. The LinkBack feature is based on the LinkBack API (which accepts the unique 'code' in an annotation ID) and text-annotator.

DRAFT

**D. Results in the Open Targets Platform.** The Europe PMC dataset is utilised in two ways: The first is to use the dataset to extract evidence for the association of targets and diseases. Co-occurrences of target and disease entities are considered evidence for the association of those entities. In detail, when a target and a disease are mentioned in the same sentence within a publication, this constitutes one piece of Europe PMC evidence for the association of that target and that disease [Figure F4 (a)]. The second involves using the dataset to provide context to the Platform entities. Users can browse the available literature for the entity of their choice through the Bibliography widget, for example all the papers linked to cystic fibrosis [Figure F4(b)]. For more details refer to article (2, 12).

### S8: Comparison with Other Tools

The outputs from the lit-OTAR framework are visualised and accessed through Europe PMC's Scilite/Annotations API and Open Targets Platform. Table T4 present comparison of various tools including SemMedDB (8), LitSense (9), PubTator (10) and PubTator Central (11) which provide similar text-mining outputs.

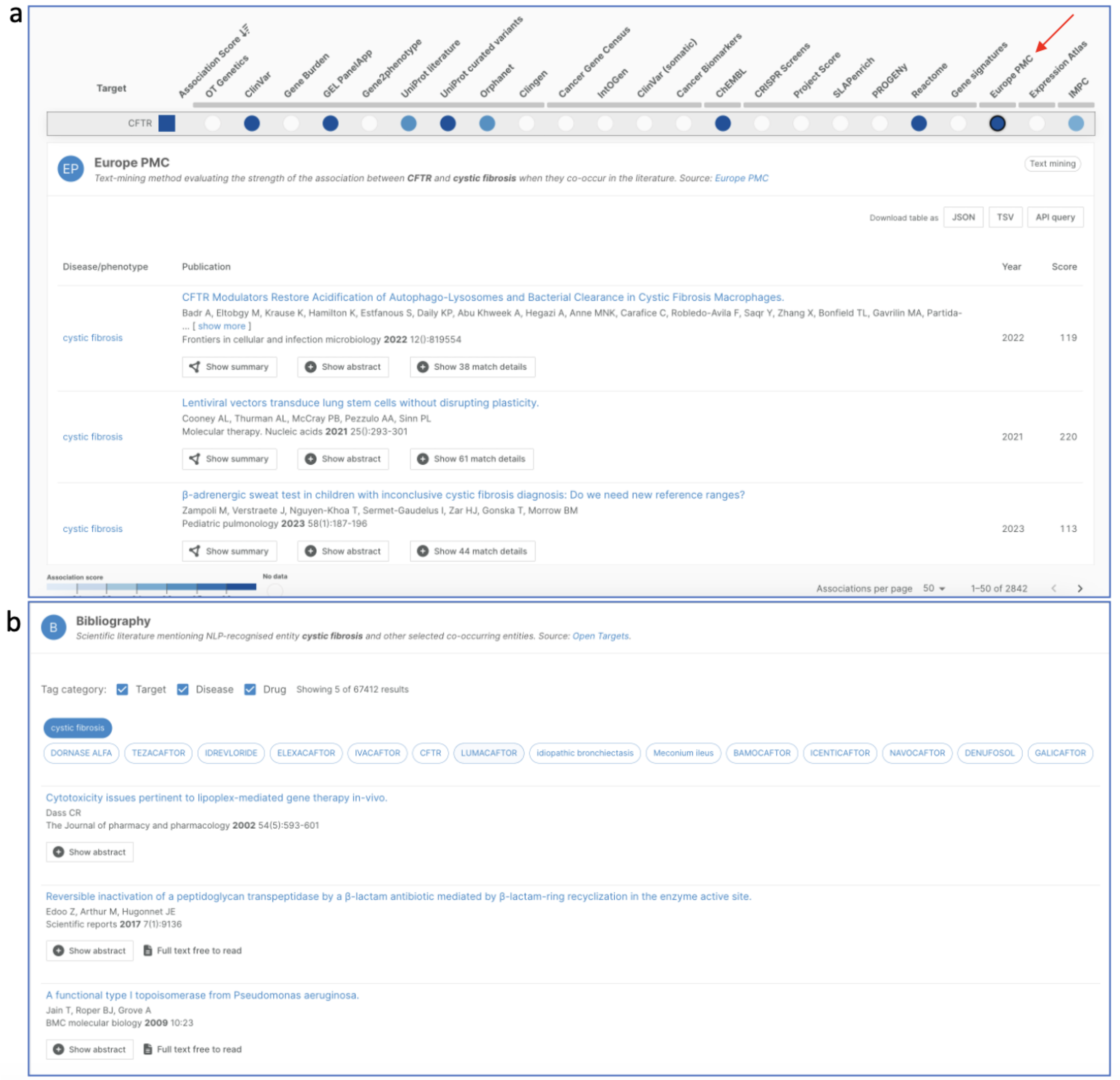


**Fig. F4.** Summary of how the lit-OTAR results are utilised and visualised in the Open Targets Platform. a. Europe PMC (red arrow) as data source for evidence of target–disease associations; b. Bibliography widget from a disease profile page.

| Feature | SciLite/Annotations API (3) | LitSense (9) | SemMedDB (8) | PubTator 3.0 (10, 11) |
|---|---|---|---|---|
| Developer | Europe PMC | NCBI | NLM (National Library of Medicine) | NCBI |
| Primary Function | Display text-mined annotations to link articles with biological data | Sentence-level retrieval of biomedical literature | Semantic predications extraction and summarization from biomedical text | text-mining, entity annotation, and relation extraction |
| Entity Types Annotated | Primarily Gene/protein names, diseases, organisms, chemicals, gene ontology terms, experimental methods, Accesion numbers, Resources. Many other entities from other providers | Genes, proteins, diseases, chemicals, mutations, species | Entities include UMLS concepts (e.g., drugs, diseases, genes, anatomy, etc.) | Genes, diseases, chemicals, variant, species, cellline |
| Relations Extracted | Gene–disease, protein–protein interactions, transcription factor–gene targets, and biological events. | None | Subject–predicate–object triples (e.g., TREATS, AFFECTS, PROCESS_OF, etc.) | 33 million relations (8.8 million unique pairs) |
| Scale of Data | Integrates multiple text-mining tools, e.g., ExTRI, IntAct, DisGeNET, PheneBank, Open Targets, and OntoGene, Metagenomics. More than 2 billion annotations. | Focused on sentence-level data | Database includes detailed structured data: citations, sentences, entities, and coreferences | 1.6 billion entity annotations (4.6 million unique identifiers) |
| Data Sources | Europe PMC articles and curated data sources (e.g., ExTRI, IntAct, Open Targets, DisGeNET) | PubMed abstracts and PMC full-text articles | PubMed abstracts; entities mapped to UMLS Metathesaurus concepts | PubMed abstracts and PMC full-text articles |
| User Interface | Highlights terms within articles and links them to external databases and tools. Exclusive API access with search | Displays relevant sentences with highlighted entities | Database schema available for querying; detailed auxiliary and semantic data accessible | Web interface and API with search |
| Update Frequency | Daily regular updates with Europe PMC content | Regular updates with PubMed and PMC content | Periodic updates; schema and data aligned with the latest biomedical literature | Weekly updates from PubMed and PMC |
| Customization | Users can select specific annotation types to display (e.g., gene-disease, protein interactions) | Users can filter results by article section or publication year | Supports custom queries on predications, coreferences, and auxiliary data | Supports semantic and relational queries with enhanced precision |
| Integration | Integrated within Europe PMC platform and connects with text-mining tools | Integrated with PubTator for entity highlighting | Can be integrated into other systems via its detailed relational schema | Integrated with NCBI resources like PubMed and PMC |
| Community | Users can upload their own data to support community | None | None | None |
| Performance | Enhanced linking of literature to biological data; focuses on annotation coverage rather than precision | Efficient for sentence-level searches | Proven semantic predication quality and flexibility in querying relationships | Relation extraction and search precision in top 20 results |

**Table T4.** Comparison of various biomedical scientific literature tools providing gene-drug-disease annotations.

DRAFT
